# Pooling quantitative MRI data: A multi-protocol study of healthy subcortical ageing

**DOI:** 10.64898/2026.01.07.698256

**Authors:** Mikhail Zubkov, Kerrin J Pine, Pierre-Louis Bazin, Puneet Talwar, Nasrin Mortazavi, Solène Dauby, Chloé Geron, Elise Beckers, Laurent Lamalle, Christophe Phillips, Fabienne Collette, Pierre Maquet, Emilie Lommers, Anneke Alkemade, Nikolaus Weiskopf, Gilles Vandewalle, Evgeniya Kirilina

## Abstract

Quantitative MRI (qMRI) measures relaxation rates, exchange rates and proton densities that reflect biophysical properties of tissue and are ideally free from protocol- and scanner-dependence. In practice, qMRI has not yet achieved this level of independence from sequence and hardware choice, and quantitative estimates often differ across sites and acquisition schemes. At the same time pooling data across different sources can be beneficial to statistical power of longitudinal, cross-sectional or case-control studies. Here we investigate how protocol and hardware differences can affect pooling data from different sources in large ultra high field (UHF) qMRI studies in the context of healthy aging. We combine the openly available ageing UHF qMRI MP2RAGEME-based dataset with two different MPM-based sets of qMRI data. We evaluate how pooling affects age dependence of qMRI parameters and investigate protocol-related biases, with a particular focus on subcortical structures. We focus the analysis, first, on replication and expansion of the reference qMRI dataset on normative aging, second, the examination of the protocol influence on the estimated qMRI values, and third, on detecting the protocol effect on the age dependence inferred from the data. We find that the age-related changes for R1 measure around 4-17% of the lifespan mean in different structures. Similarly, age-related R2* variation in different structures constitutes around 6-30%. Subcortical structure volume change is on the order of 5-27%. We further observe larger relative difference between protocols for R1 and volume, while R2* remains more consistent for most regions. We show how pooling the UHF qMRI data from different sites and collected with different quantitative protocols can be both detrimental and beneficial for the analysis outcomes.

## Introduction

Ultra-high field (UHF, i.e. at 7T and above) quantitative MRI (qMRI) with its ability to provide ultra-high resolution quantitative maps of brain anatomy, is progressively entering both the clinical and research applications. Recent developments in parallel radio frequency (RF) transmission techniques (pTx), have mitigated biases caused by transmit field inhomogeneity inherent to UHF MRI (Webb & Collins, 2010), substantially advancing its applications. Owing to these advances, elusive changes in smaller subcortical nuclei as well as details of cortical laminar structures can now be detected non-invasively (Miletić et al., 2022; Trampel et al., 2019). This enables neuroscientists and clinicians to better characterize their anatomy in healthy population, and to identify subtle alterations associated with brain disorders and neurodegenerative diseases (Forstmann et al., 2017; Keuken et al., 2018).

qMRI measures absolute spin ensemble characteristics (i.e. relaxation rates, exchange rates and proton densities (PD)) that reflects biophysical properties of tissue and aim to minimize protocol- and scanner-dependence (Weiskopf et al., 2021). By linking these qMRI parameters to tissue microstructure via biophysical models, qMRI provides insights into the tissue microarchitecture beyond the nominal resolution of acquired data. In particular, the longitudinal relaxation rates (R1) and the effective transverse relaxation rate (R2*) have been shown to yield unique information about the tissue macromolecular and iron content (Mackay et al., 1994; Metere & Möller, 2016; Sereno et al., 2013; Stüber et al., 2014). The combination of UHF and qMRI is therefore very promising for enabling true *in vivo* histology MRI (hMRI(Weiskopf et al., 2021)) of small brain structures, such as cortical layers and subcortical nuclei. This creates opportunity for a new generation of early-stage, non-invasive MRI biomarkers of neurodegeneration, as small subcortical nuclei are often affected in early pre-symptomatic disease stages. Realizing this potential, however, requires normative populational datasets that capture the full variability of anatomy and microstructure across the healthy adult lifespan. Such studies are pivotal for defining quantitative reference values, discriminating between normal aging and pathology, and setting the basis of clinical diagnostics.

Several approaches for multiparametric qMRI have been developed. Most frequently one of the following two is used. Multi-echo magnetization-prepared rapid gradient echo (MPRAGEME or MP2RAGEME), based on application of inversion pulse followed by rapid gradient echo readouts, simultaneously yields (Caan et al., 2019) the quantitative maps of R1 and R2*. Alternatively, variable excitation flip angle protocols, such as multiparameter mapping (MPM), also provide (Weiskopf et al., 2013) R1 and R2* maps. Ideally, these qMRI techniques should yield consistent results across protocols and scan sites, while preserving the sensitivity to microstructural and gross anatomical details of individual participants.

In practice, however, UHF qMRI has not yet achieved full independence from protocol and hardware(Bauer et al., 2010; Lee et al., 2019). In addition to technical reasons, remaining variation is due to the fundamental challenge to consistently capture complex tissue compositions (e.g. multiple compartments), which cannot be perfectly described by a single parameter and are the objective of biophysical modeling and in-vivo histology approaches (Weiskopf et al., 2021). Nevertheless, qMRI studies were able to infer the changes of brain microstructure in ageing (Carradus et al., 2020; Keuken et al., 2017; Miletić et al., 2022; Piredda et al., 2023), in depressive disorder (Heij et al., 2024), in multiple sclerosis (Al-Radaideh et al., 2015; Lommers et al., 2021), and in COVID-19 (Rua et al., 2024). Large cohort studies of aging have consistently identified inverted U-shape age dependence of R1, R2*, and brain volume across multiple brain regions, reflecting protracted brain plasticity up to late adulthood followed by age-related decline. Disease-oriented studies usually rely on normative values by monitoring the parameter deviations in selected brain structures as indicator of pathology. To create such normative large representative datasets, pooling data across different sites and protocols is highly beneficial, as it increases statistical power of longitudinal, cross-sectional or case-control studies. However, the lack of unified mapping protocols in UHF qMRI and the consequential lack of consensus on normal values of tissue parameters reduces the prognostic utility of the method, prohibiting introduction of cutoff values for disorder diagnostics as inter-individual (Callaghan et al., 2014; Miletić et al., 2022) and between protocol differences cannot be properly accounted for.

Encouragingly, recent studies (Leutritz et al., 2020; Pine et al., 2024; Rua et al., 2020) have demonstrated that when site differences are taken into consideration and protocol consistency is ensured, the qMRI estimates become generalizable at the participant level. In other words, scanning the same person across different sites with the same protocol yields similar outputs. Building on this finding, here we investigate how protocol and hardware differences can affect pooling data of different individuals in large qMRI studies in the context of healthy aging. We combine the openly available ageing UHF qMRI reference dataset (Alkemade et al., 2020) with heterogeneous (in terms of the protocol in use) new data. Specifically, we evaluate how pooling datasets from different sites affects estimates of age dependence of qMRI parameters. We investigate protocol-related biases, with a particular focus on subcortical structures, whose reliable delineation requires UHF MRI due to their small size and segmentation challenges.

## Methods

### Participant cohort

The sample used in this study comprises three cohorts of healthy control participants with ages covering the human adult lifespan, from 18 to 80 years old, and originates from three different sources, referred here as AHEAD, sTx-MPM and pTx-MPM. The AHEAD study is sourced from a publicly available dataset (Alkemade et al., 2020) acquired with the MP2RAGEME sequence on a Philips Achieva 7T (Philips Healthcare, Eindhoven, Netherlands) scanner at the University of Amsterdam. AHEAD contains 105 participants (60 female, 45 male) aged from 19 to 80 years. The other two cohorts’ data sources are the studies conducted at the GIGA-CRC Human Imaging unit of the Université de Liège on a 7T Siemens Terra (Siemens Healthineers, Erlangen, Germany) scanner and are not yet publicly available. This new data consists of two parts making use of different versions of the MPM approach: one labelled as sTx-MPM and another labelled as pTx-MPM (see Table 1). Both versions employ a multi-echo (ME) magnetization transfer (MT) enabled Fast Low Angle Shot (FLASH)-based MRI protocol, but with variations in scan parameters and hardware (see Table 2 and a detailed description below). AHEAD measurements serve as a reference data for both MPM datasets due to larger cohort.

**Table 1.**
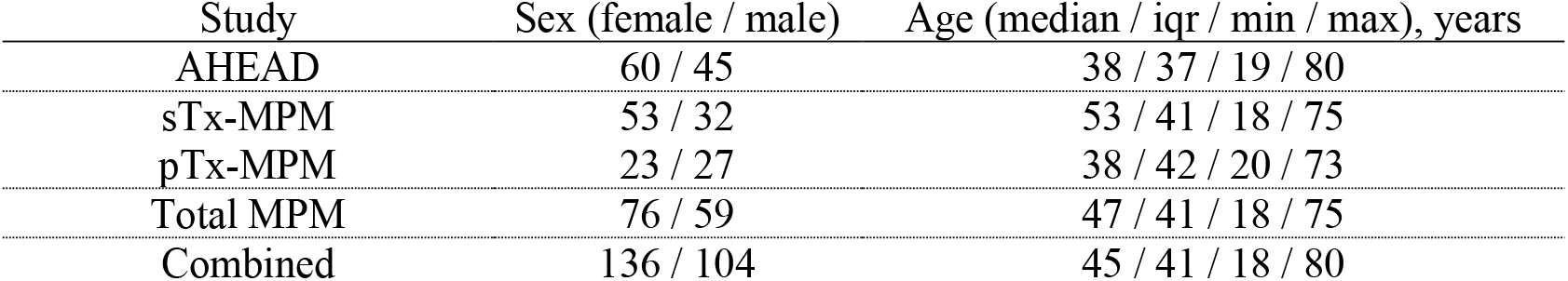
Age and sex distribution of the subsamples comprising the study cohort. Age column contains median values, interquartile range (25^th^ to 75^th^ percentile), minimum and maximum age values.

**Table 2.** Major features of the reference and new acquisitions. The full parameters for MT-FLASH-ME acquisition sequences, as well as for Spin Echo/Stimulated Echo Echo-Planar Imaging (SE/STE-EPI) based B1^+^ field mapping and Actual Flip Angle (AFI) sequence are provided in the Data acquisition section.

|  | AHEAD | sTx-MPM | pTx-MPM |
| --- | --- | --- | --- |
| Scanner | Philips Achieva 7T | 7T Siemens Terra | 7T Siemens Terra |
| Coil | 1Tx/32Rx Nova Coil | 1Tx/32Rx Nova Coil | 8Tx/32Rx Nova Coil |
| Sequence | MP2RAGEME | MT-FLASH-ME | MT-FLASH-ME |
| Number of echoes | 4 | 4 and 6 | 4 and 6 |
| Resolution | <0.7 mm | 0.6 mm | 0.6 mm |
| B1<sup>+</sup> mapping | none | SE/STE-EPI | AFI |
| Metrics extracted | R1, R2*, $\chi$ | R1, R2*, PD, MTsat | R1, R2*, PD, MTsat |
| qMRI tool | nighres, tgv-qsm | hMRI-toolbox 0.6.1 | hMRI-toolbox 0.6.1 |

Each study was approved by the corresponding ethics committee. Written informed consent was obtained from all participants for being included in the study. The exclusion criteria for the participants were history of major neurologic/psychiatric diseases or stroke; recent history of depression/anxiety; sleep disorders; medication affecting the central nervous system; smoking, excessive alcohol (>14 units/week) or caffeine (>5 cups/day) consumption; night shift work in the past 6 months; BMI ≤18 and ≥29 (for older individuals) and ≥25 (for younger individuals). The separate statistics for the studies are presented in Table 1. The total MPM sample comprised 135 participants (76 female, 59 male) aged from 18 to 75 years. As described in detail below the data analysis employed the pooled dataset (combining both MPM and the AHEAD data), producing a combined dataset counting 240 participants with further details provided in Table 1.

### Data acquisition

#### AHEAD

The full acquisition protocol details of the AHEAD data are described in the original publication (Alkemade et al., 2020). Briefly, the study made use of a single transmit 1-Tx/32-Rx RF head coil (Nova Medical, Wilmington, MA, USA) and employed an MP2RAGEME acquisition (Caan et al., 2019) with two inversion times (TI1/TI2 670.0/3675.4 ms) and four unequally spaced echoes (echo time (TE) of 3.0/11.5/19/28.5 ms). The scan covered the Field of View (FoV) of 205×205×164 mm^3^ with a 0.64×0.64×0.7 mm^3^ nominal resolution. The acquisition was accelerated using parallel imaging with the SENSitivity Encoding (SENSE) technique (Pruessmann et al., 1999) with an acceleration factor of two in one phase encoding direction. No transmit B1^+^ field inhomogeneity correction was performed.

#### sTx-MPM

Acquisition of the data labelled sTx-MPM employed scanning with a 1-Tx/32-Rx head coil (Nova Medical, Wilmington, MA, USA) and an MPM protocol (Vaculčiaková et al., 2022; Weiskopf et al., 2013) comprising three (T1-weighted, PD-weighted, MT-weighted) whole-brain 3D FLASH-based multi-echo sequences covering the FoV of 256×218×173 mm^3^ with isotropic resolution of 0.6 mm (repetition time (TR) 19.5 ms, flip angles (FA) PDw/MTw/T1w 5/5/20°, six equispaced echoes with TE ranging from 2.3 to 14.2 ms for PDw and T1w and four equispaced echoes with TE ranging from 2.30 ms to 9.44 ms for MTw (the MT-weighting was achieved by applying a 4 ms, 140° Gaussian-shaped pulse, 2 kHz off-resonance), parallel imaging using GeneRalized Autocalibrating Partial Parallel Acquisition (GRAPPA) approach (Griswold et al., 2002) with acceleration factor of two in both phase encoding directions, and 3D spin-echo/stimulated-echo echo planar imaging (SE-STE-EPI) based transmit B1^+^ field mapping for transmit field inhomogeneity correction (Jiru & Klose, 2006). This field mapping protocol was acquired at an isotropic 4 mm resolution (TR 500 ms, TE 43.18 ms, and 15 gradually decreasing nominal pulse flip angles from 165° to 60°, GRAPPA acceleration in both phase encoding directions with acceleration factor R=2 in each). A dual-echo gradient echo sequence with 3 mm isotropic resolution (TR 677 ms, FA 39°, TE 5 ms and 6.02 ms) was acquired for B_0_ mapping employed in EPI distortion correction.

#### pTx-MPM

Acquisition of the data labelled pTx-MPM adhered to a protocol making use of the parallel transmission using a 8-Tx/32-Rx RF head coil (Nova Medical, Wilmington, MA, USA) and an MPM protocol comprising three (T1-weighted, PD-weighted, MT-weighted) whole-brain 3D FLASH-based multi-echo sequences covering the FoV of 250×218×173 mm^3^ with isotropic resolution of 0.6 mm (TR 22.4 ms, FA PDw/MTw/T1w 7/7/22°, six equispaced echoes with TE ranging from 3.0 ms to 15.6 ms for PDw and T1w and 4 equispaced echoes with TE ranging from 3.00 ms to 10.56 ms for MTw (MT weighting achieved by applying a 4 ms, 130° Gaussian-shaped pulse, 3 kHz off-resonance), Controlled Aliasing In Parallel Imaging Resulting IN Higher Acceleration (CAIPIRINHA) acceleration (Breuer et al., 2006) in two phase encoding directions with acceleration factor R=2 in each, kT-points pulses (Pine et al., 2023, p. 500) for RF homogeneity improvement and AFI based B1^+^ mapping for transmit field correction (Yarnykh, 2007). The field mapping protocol was acquired at an isotropic 4 mm resolution (two TR values of 25 ms and 125 ms, TE 2.12 ms, nominal pulse flip angle of 55°, GRAPPA acceleration in one phase encoding direction with an acceleration factor R=2). Two field mappings were performed per protocol to account for different B1^+^ distributions when using the kT-points excitation pulses and the Gaussian-shaped MT-pulses.

To test if different B1^+^ field mapping procedures in sTx-MPM (SE-STE-EPI) and pTx-MPM (AFI) could be a source of error, particularly for such B1^+^ sensitive measurements as R1 mapping, three participants from the pTx-MPM cohort also had the SE-STE-EPI based field mapping with the parameters matched to the field mapping from the sTx-MPM added to their scan protocols.

### Data processing pipelines

The reference AHEAD data processing pipeline for quantitative parameter extraction is described in the original publication (Miletić et al., 2022). Briefly, it comprises the LCPCA-based denoising (Bazin et al., 2019), standard MP2RAGE calculation of the T1 relaxation time (Marques et al., 2010), single-exponential fitting of the multi-echo data for the R2* estimation, and TGV-QSM (Langkammer et al., 2015) for QSM. The output quantitative maps (R1, R2*, and χ) were then used in the Multi-contrast Anatomical Subcortical Structures Parcellation (MASSP) (Bazin et al., 2020) Bayesian segmentation algorithm based on anatomically-derived priors and contrast-conditional posteriors yielding masks of 14 subcortical structures (left/right averaged).

The newly acquired data from sTx-MPM and pTx-MPM studies was processed through two similar pipelines, which included LCPCA-based denoising of the anatomical magnitude images, hMRI-toolbox (www.hmri.info, version 0.6.1) based quantitative parameter maps calculation (Tabelow et al., 2019), employing corresponding incomplete spoiling correction, calculated for each pulse sequence, and two-iteration weighted least squares R2* fitting. The final processing step comprised MASSP-based segmentation, as the latter has demonstrated reliable and reproducible segmentation in both AHEAD data and dedicated reproducibility study (Pine et al., 2024).

The difference in processing the sTx-MPM and the pTx-MPM data was, first, in different models for B1^+^ maps calculation, and second in employing different B1^+^ inhomogeneity corrections for the MT-pulses, the latter of least importance due to MTsat maps being unused in this study. The qMRI mapping yielded the PD, R1, R2*, and MTsat maps, with the first three having been used for the MASSP parcellation (and the MTsat excluded from further processing), that provided segmentation of the same 14 subcortical structures (left/right hemisphere averaged, see Figure 1 for details) as in the AHEAD data. Aside from the difference in acquisition protocols, a crucial data-wise difference between AHEAD and MPM data is that for MPM scans only magnitude data were exported, thus no QSM data was reconstructed neither for the sTx-MPM nor pTx-MPM protocols. On the other hand, recovering the PD metric allowed to incorporate it into the segmentation procedure in place of QSM.

**Figure 1.**
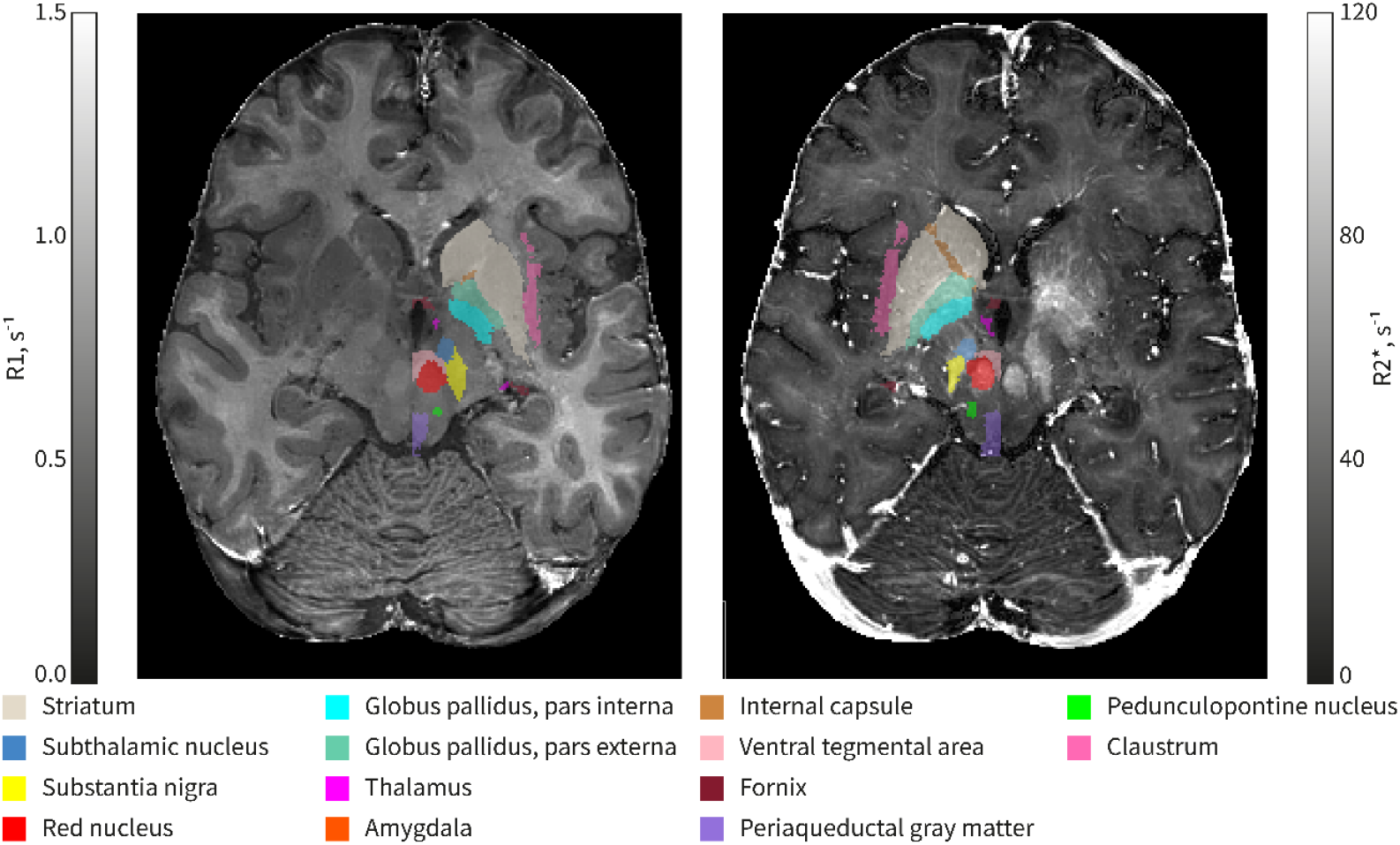
Segmentation example of individual dataset (pTx-MPM, f, 46 years) showing a set of investigated 14 subcortical structures overlaid on the pTx-MPM based R1 parametric map (left) and R2* parametric map (right). Segmentation in only one hemisphere is shown for each map.

Two additional analyses were performed in order to estimate the biases resulting from different B_1_^+^ mapping methods and different B_1_^+^ field settings employed in sTx and pTx acquisitions. To estimate the effect of different mapping on qMRI parameters, the B_1_^+^ field mapping data for three participants having both types (AFI and SE-STE-EPI) of field mapping was used to calculate the relative flip angle maps (in percent units of nominal flip angle) and was co-registered to anatomical data using the SPM12 tools via the hMRI toolbox. For these participants, joint histogram of relative flip angle values estimated by two methods was plotted, and linear model assessing the ratio between the two measurement methods was fit to the data. To estimate the effect of different B_1_^+^settings in sTx and pTx acquisitions the corresponding relative flip angle maps for all other participants were used in conjunction with MASSP-based segmentations to calculate the mean relative flip angle and the coefficient of variation (CoV) of the relative flip angle across the 14 subcortical structures.

### Quality control

The data was visually checked for presence of gross movement, artifacts and segmentation failures. Potential errors in segmentation and qMRI maps calculation were also assessed by plotting the ratio of non-unique (i.e., values different at the digital number format precision) R2* values per segmented structure volume for all ROIs and participants. Due to continuous distribution of R2* values in noisy data, all voxels should possess unique values, and the number of non-unique values should be zero. Systematic errors, e.g., convergence of the fitting procedure to boundary values may produce duplicated values, which can be identified via calculating of the ratio of non-unique values to the ROI volume. Another source of systematic error is the low-voxel count ROIs: these can be generated due to the segmentation errors, and may produce outlier values, while being undetected by the non-unique value count procedure (e.g., a single-voxel ROI has no duplicated values, but is likely to produce outliers). Shannon entropy (Shannon, 1948) allows assessing the amount of information the ROI can provide, and favors larger ROIs with non-repeated values. Practically, entropy can be easily thresholded, e.g., excluding values less than 3 bits, thus effectively removing areas less than 8 voxels (particularly, single-voxel segmentations with zero entropy), as well as larger areas with sufficiently frequent value repetition – a common sign of boundary value convergence.

### Statistics

Median values for each of the calculated quantitative parameters were extracted for the 14 subcortical structures (average of left and right ROIs) with the ventricular system excluded from further analysis. The statistical analysis described below was carried out separately for each of the three different qMRI metrics (R1, R2*, and ROI volume) using the combined cohort comprising 240 healthy participants.

The statistical analysis focused on three goals. First, a replication and expansion of the reference qMRI dataset on normative aging, i.e., the inference of life-span trajectory of subcortical anatomy and microstructure in healthy aging, as measured with UHF qMRI. Second, the examination of the protocol influence on the estimated qMRI values (i.e., the detection of protocol-related absolute biases in age-corrected median values of qMRI parameters), and third, assessing if acquisition protocols impact the estimates of the age dependence of qMRI parameters (i.e., the interaction between protocol effect and estimated aging trajectory). To achieve these three objectives, a linear model was built in R 4.4.0 for each of the median qMRI metrics with age, age^2^, and sex as regressors (with the second power of age included following the results in (Carradus et al., 2020; Miletić et al., 2022; Piredda et al., 2023)). In order to assess the impact of different protocols on the estimated median values, an additional categorical parameter related to the protocol was incorporated into the models with three categories indicating either sTx-MPM, pTx-MPM, or AHEAD data. Furthermore, interaction between the protocol-related term and the age-related variables were considered.

The presence of age and protocol dependence was assessed via various F-tests over a set of models. The set of F-tests to detect age dependence (goal 1) compared models with age, age^2^, sex, and protocol type variables against models with only sex, and protocol type variables. The bias detection (goal 2) was conducted via an F-test of the model with age, age^2^, sex, and protocol type variables against the model without the protocol type variable. To detect the influence of protocol selection on age dependence (goal 3) models with age, age^2^, sex, and protocol type variables were tested against models with added interactions between age, age^2^ and protocol type variables. Each of the qMRI parameters was treated as a distinct measurement and multiple comparison correction was applied across different subcortical structures separately for each extracted qMRI metric and for each type of test. The effects with F-test p-values lower than 0.05 threshold (Benjamini–Hochberg false discovery rate-corrected (Benjamini & Hochberg, 1995)) were deemed significant.

To further assess the influence of the protocol on the estimated qMRI parameters, first, the age trajectories were used to calculate the variation of the qMRI parameters with age as the difference of the maximum and minimum values within the studied age range. These variation estimates were then used to assess how different were the protocol-specific quantitative values from the reference AHEAD protocol as compared to the age-related variation of each quantitative parameter estimated using sTx-MPM and pTx-MPM datasets.

Age trajectories of R1 reflecting the processes of myelination in young adulthood and demyelination in older age were used to calculate the age where myelination reached its maximum for each segmented brain region.

## Results

### Acquisition and segmentation

The combination of the sTx-MPM and pTx-MPM datasets (see Table 1) resulted in a 135-participant dataset with age distribution similar between all three (sTx-MPM, pTx-MPM and AHEAD) cohorts (Figure 2).

**Figure 2.**
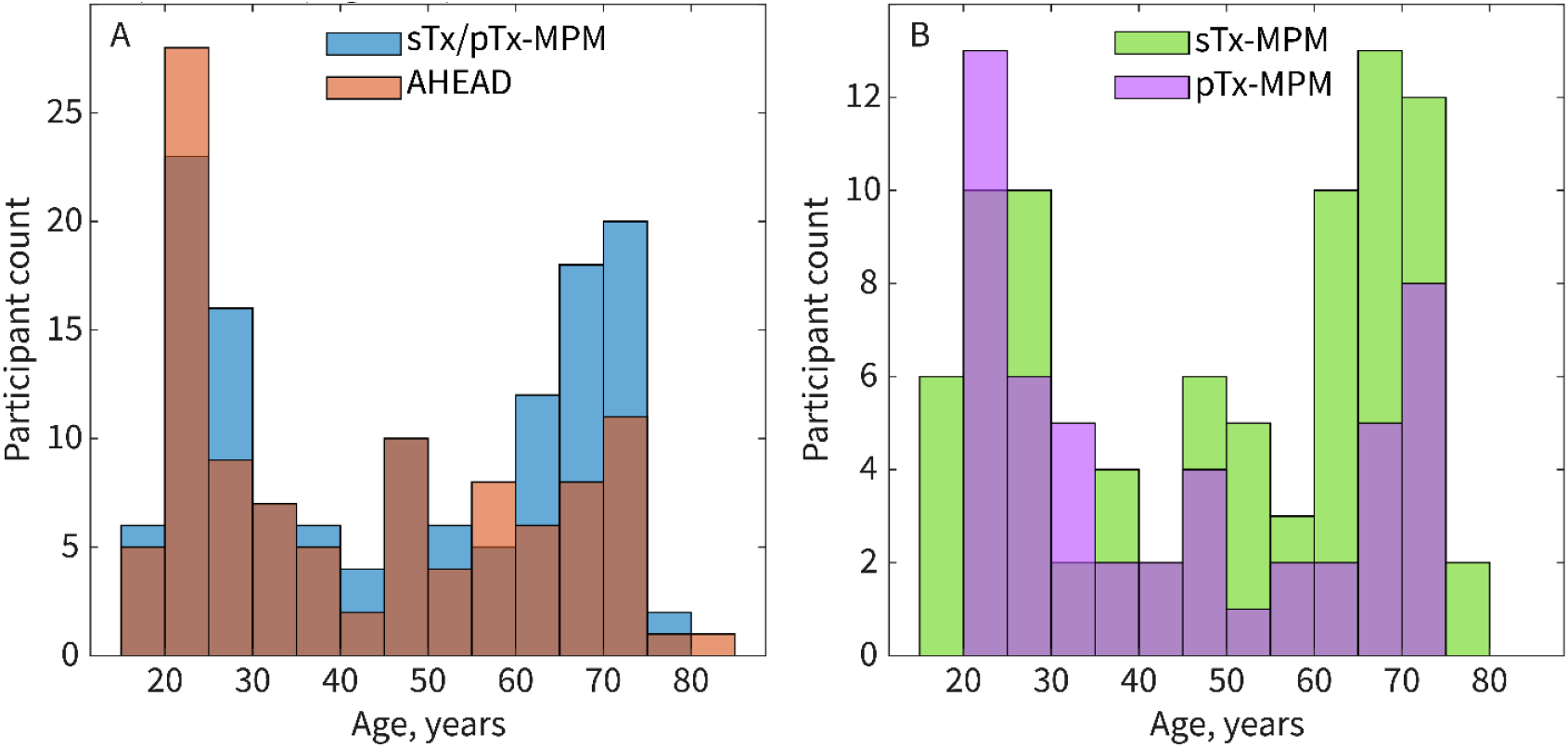
Overlaid histograms of age distribution in the participant cohort. A. Age distribution in the AHEAD (red) and MPM (sTx-MPM and pTx-MPM), blue) datasets. The similarity of the age profiles justifies employing similar approaches to data analysis as in (Miletić et al., 2022) and allows comparing age dependence. B. Age distribution in the two parts of the MPM dataset (green for sTx-MPM and purple for pTx-MPM). Again, the distribution similarity allows pooling without risk of causing age biases by particular cohort subset.

High quality, 0.6 mm resolution, co-aligned maps of qMRI parameters R2*, PD, R1 were obtained in all participants of the combined studies. The resolution of the parametric maps and the contrast in cortical and subcortical anatomical structures allowed MASSP parcellations (Bazin et al., 2020) of the subcortical structures highlighted in Figure 1 in each participant. The quality assurance procedure confirmed that all qMRI parameter estimations and segmentation performed well (Figure 3). Particularly, no regions exhibited higher than 0.3% of non-unique values per ROI volume. No regions with Shannon entropy (Shannon, 1948) values less than 6 bits were observed, thus allowing to include all of the segmented regions into analysis and suggesting substituting χ with PD measures did not dramatically degrade MASSP segmentation quality.

**Figure 3.**
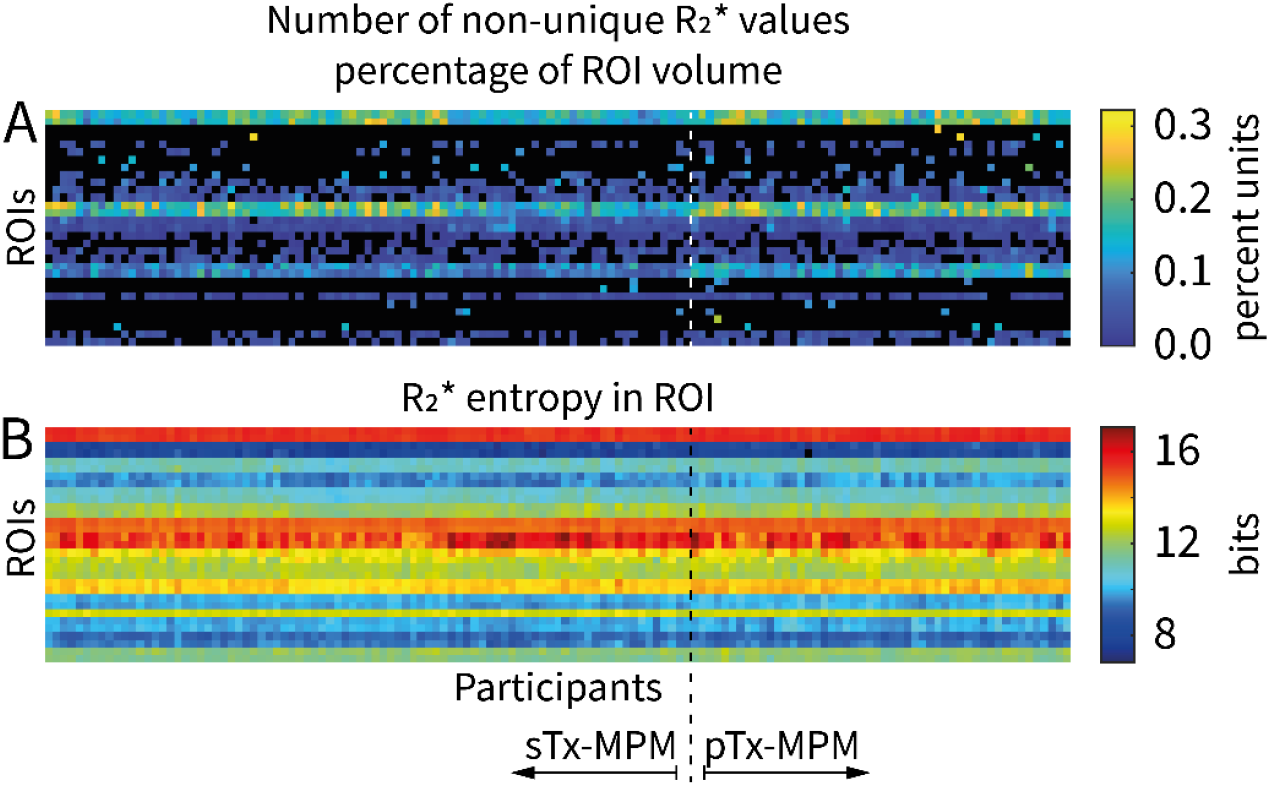
Quality assurance results for the newly acquired data. Note, that the number of ROI is higher than the one described in the Methods section due to the quality metrics being extracted prior to averaging left and right structures. A. The overall low percentage (well below 1%) of non-unique values in all ROIs indicates lack of convergence of qMRI metric calculation to boundary values. B. High entropy (above 6 bits) in all the ROIs indicates absence of low-voxel count ROIs and absence of undesired convergence to boundary values.

Data and qMRI parameter age dependence extracted from a number of representative ROIs (Striatum, Globus Pallidus and Thalamus) are provided in Figure 4. Data for all 14 segmented regions are provided in Supplementary Figures S1 to S6.

**Figure 4.**
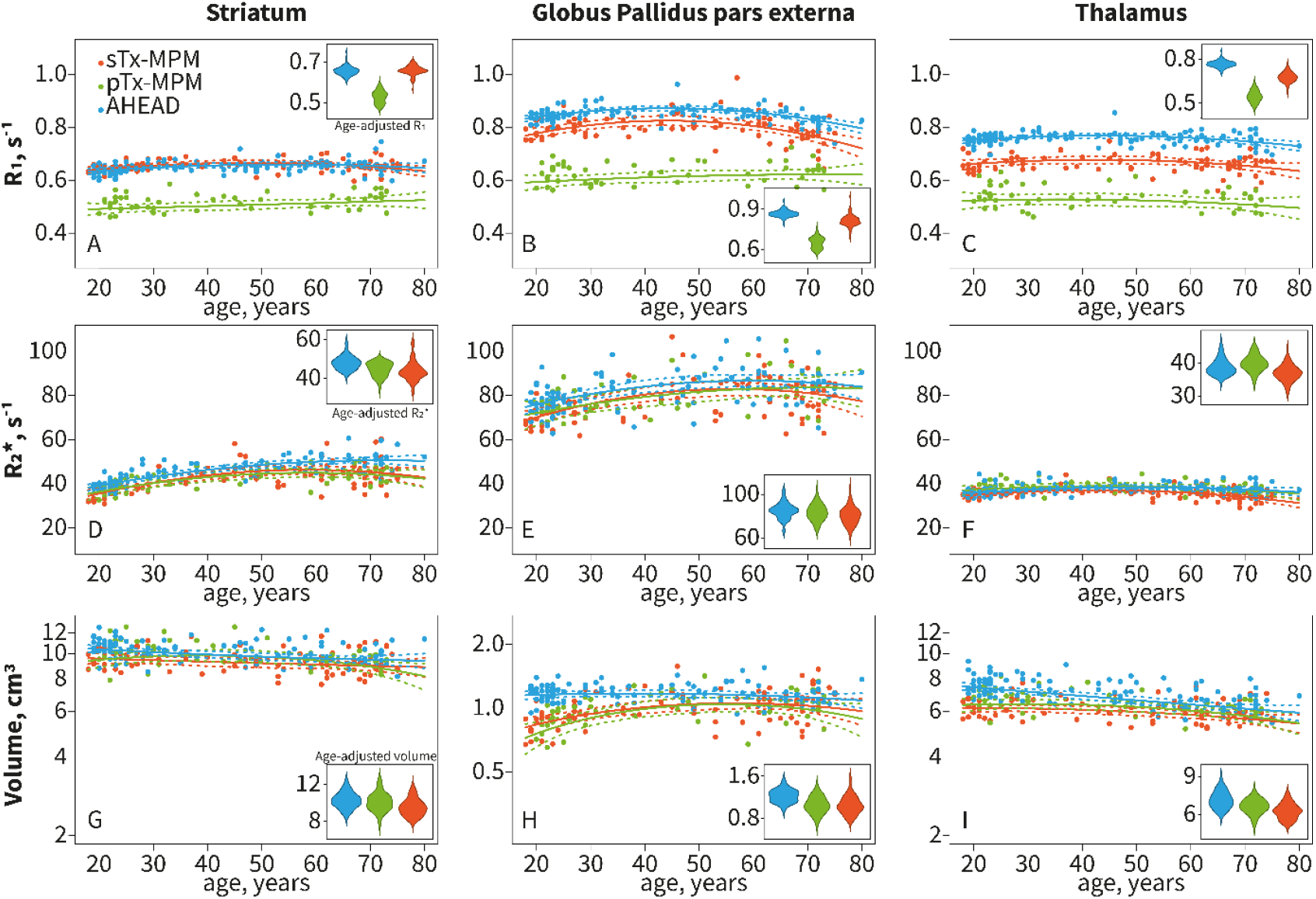
Examples of R1 (A, B, C), R2* (D, E, F) and ROI volume (G, H, I) age dependence in Striatum (A, D, G), Globus Pallidus pars externa (B, E, H), and Thalamus (C, F, I). Each graph also shows models that were fit for individual protocols (not used in statistical analyses). Insets show age-adjusted data distribution across different protocols, where all participant data has been adjusted to median cohort age (45 years). Note the volume plots use different logarithmical scales across different ROIs.

### Statistics for the combined dataset

We start with the description of the age-related changes in the qMRI metrics. Figure 5 provides a representation of effect size for the effect of interest in all 14 ROI. The generalized linear models showed significant (p<0.05, FDR-corrected) combined age and age^2^ dependence in R_1_, R2* and ROI volume for all of the segmented subcortical structures, except for the Periaqueductal Gray volume (see significance indicated via * in Figure 5).

**Figure 5.**
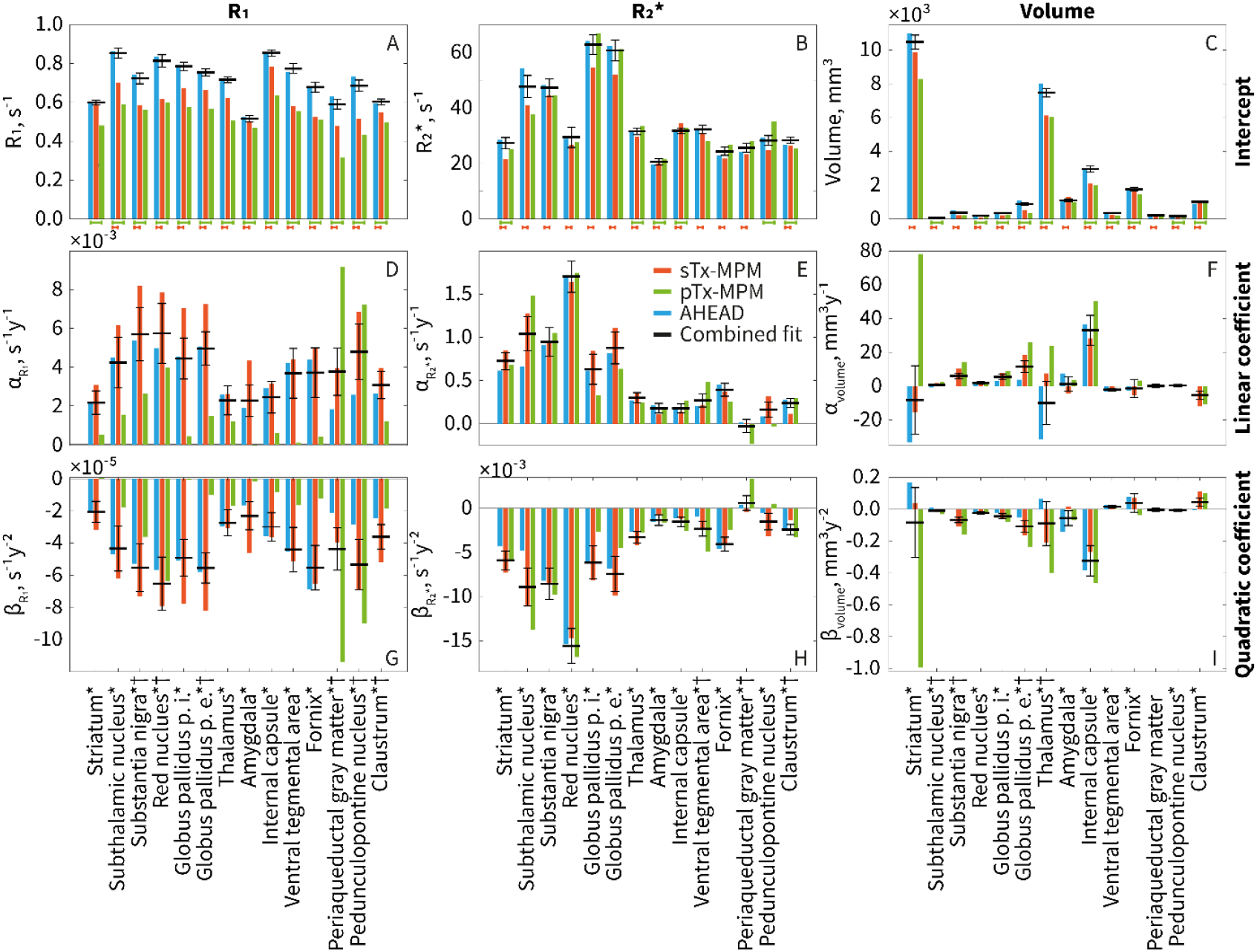
Age-dependence linear model (metric ∼ intercept + α*age + β*age^2^) coefficients for R1 (A, D, G), R2* (B, E, H), and ROI volume (C, F, I). Colored columns show effect size for per-dataset fits (blue for AHEAD data, red for sTx-MPM, green for pTx-MPM), black bars indicate effect sizes for a combined dataset fit. Top row (A, B, C) shows the model intercept, middle row (D, E, F) shows the model linear coefficient, bottom row (G, H, I) shows the model quadratic coefficient. Asterisk (*) following the ROI name indicates statistical significance of age dependence in the combined dataset. Red bar under intercept graphs (A, B, C) indicates statistically significant differences between the sTx-MPM data and AHEAD data, green bar under intercept graphs indicates statistically significant differences between the pTx-MPM data and AHEAD data. Dagger (†) indicates statistically significant interaction between protocol-related categorical variable and age-related variables.

Next, we focus on the differences between protocols (see color-coded significance across Figure 5). When comparing the MPM data with the AHEAD data, the models showed significant differences of R1 values measured with the pTx-MPM protocol in all the ROIs, and with the sTx-MPM protocol in all ROIs except Striatum (see Figure 5 A for the effect sizes). Significant differences between the AHEAD protocol and the pTx-MPM protocol were detected in R2* in Striatum, Thalamus, Amygdala, Internal Capsule, Pedunculopontine Nucleus, and Claustrum. Differences in R2* between the AHEAD and the sTx-MPM protocol were found in all ROIs except Pedunculopontine Nucleus (see Figure 5 B for the effect sizes). Furthermore, differences in ROI volume measured with the pTx-MPM protocol, compared to the ones calculated in the AHEAD data were found in all ROIs except Striatum, Substantia Nigra, Amygdala, and Periaqueductal Gray, and with the sTx-MPM protocol – in all ROIs (see Figure 5 C for the effect sizes).

Finally, we consider the influence of the protocol on the age dependence (see significance indicated via † in Figure 5). Significant interactions between age-related variables and protocol were detected for R1 in Substantia Nigra, Red Nucleus, Globus Pallidus (pars externa) Periaqueductal Gray, Pedunculopontine Nucleus, and Claustrum; interactions for R2* measurements were detected in Ventral Tegmental Area, Periaqueductal Gray, and Claustrum; and interactions in volume measurements were detected in Subthalamic Nucleus, Substantia Nigra, Globus Pallidus pars externa, and Thalamus.

The expected age-related change in quantitative metrics varied across different ROI between 3.8% (0.025 s^-1^) and 16.8% (0.119 s^-1^) of the mean value for R_1_ (0.65 s^-1^ and 0.71 s^-1^ respectively), between 6.4% (2.06 s^-1^) and 29.7% (20.96 s^-1^) for the R_2_^*^ (mean values of 32.29 s^-1^ and 70.55 s^-1^ respectively) and between 4.9% (10.7 mm^3^) and 26.5% (268.1 mm^3^) for the region volume (mean values of 220.8 mm^3^ and 1010.2 mm^3^ respectively). At the same time (Figure 6) the mean absolute protocol-related difference from the AHEAD data for the sTx-MPM was 140% of the expected age-wide variation for R1, 70% for R2*, and 161% for the volume. For the pTx-MPM the values were 413% of the expected age-wide variation for R1, 36% for R2*, and 123% for the volume. When referenced to the expected mean value of the quantitative metric (instead of the expected age-wide variation), the sTx-MPM protocols exhibited on average a 9.5% lower R_1_ values, a 6.6% lower R_2_^*^, and 17% lower volume. For the pTx-MPM the R_1_ values were on average 27% lower than in the AHEAD data, the R_2_^*^ values were 0.5% higher, and regional volume was 12.7% lower. The ages of maximal R1 calculated from the age trajectories are presented in the Supplementary Figure S7 and exhibit levels of error rendering them hardly practical.

**Figure 6.**
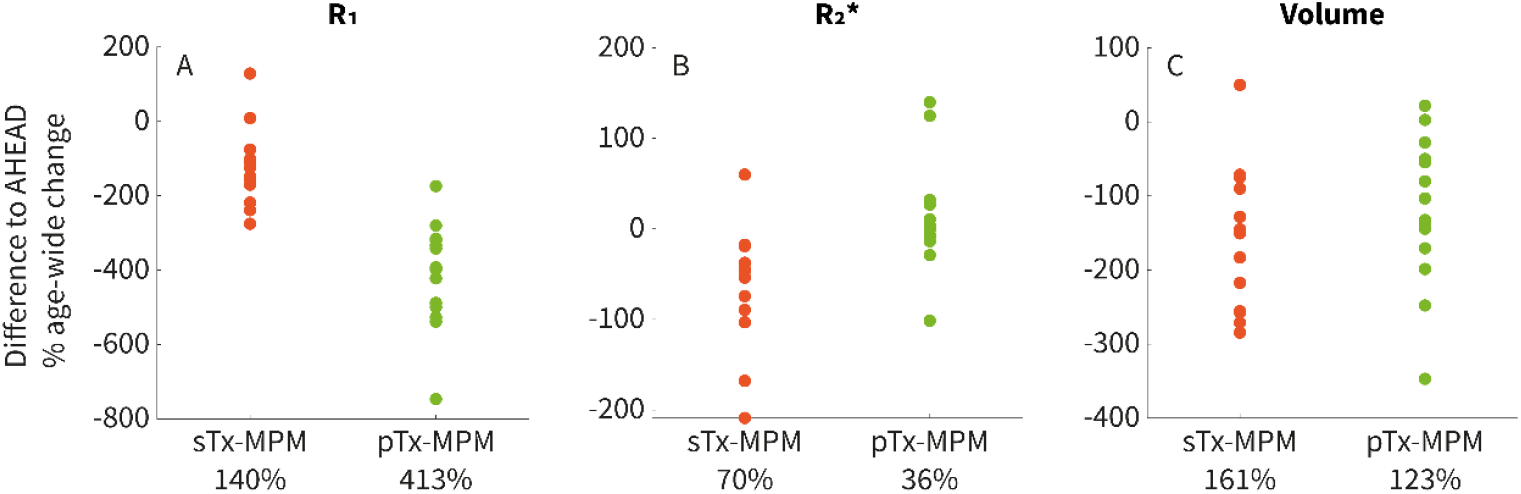
The effect of protocol selection on the measured protocol differences in quantitative value normalized to the age-related change in the metric, shown for 14 subcortical structures (dots). A – pTx-MPM (green) exhibits lower R_1_ values with mean absolute difference of 413%, while sTx-MPM (red) demonstrates a difference of 140% of the expected age-related change (significant with p<10^-5^). B – both sTx-MPM and pTx-MPM exhibit average difference in R_2_^*^ lower than the expected age-related change, 70% and 36% correspondingly (difference significant with p<0.001). C – both sTx-MPM and pTx-MPM show comparable differences against AHEAD data (161% and 123% of the expected age-related change, significant difference only with p<0.05).

### B_1_^+^ field measurements

The analysis of potential bias resulting from different B_1_^+^ mapping procedure and different B_1_^+^ settings in sTx and pTx protocols are shown in Figure 7. Comparison of different B_1_^+^ mapping techniques (Figure 7 C), indicated that in three subjects where both AFI and SE-STE-EPI based field measurements were performed, the resulting field maps matched closely. Particularly, linear models with intercept fixed at zero (i.e., assuming both methods showing zero field when no field is present), demonstrated a slope of 1.002 with adjusted r-square of 0.9657.

**Figure 7.**
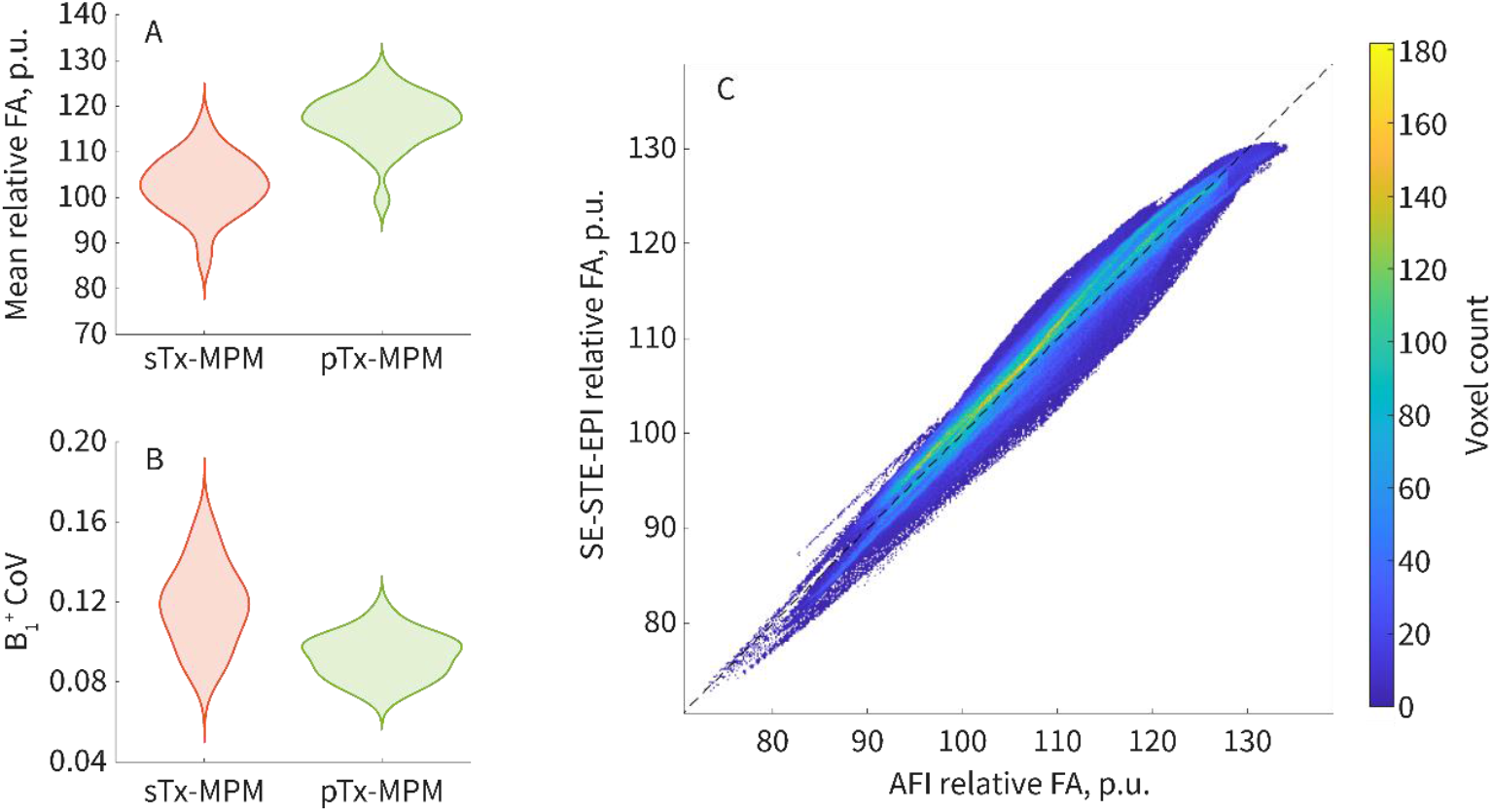
A – Distributions of the estimated B_1_ ^+^ values in the MPM cohort (in percent units relative to value corresponding to the nominal flip angle) averaged across all the segmented subcortical structures. B – Distribution of the coefficient of variation (CoV) of the estimated B_1_^+^ values across all subcortical structures. Graph A demonstrates the pTx-MPM protocol to produce higher B_1_ ^+^ values, which also vary less from subject to subject. Graph B demonstrates the pTx-MPM protocol to produce more uniform B_1_ ^+^ distribution with less homogeneity variation across subjects. C – joint histogram of B_1_ ^+^ estimated consecutively in the same subjects with SE-STE-EPI and AFI methods. The histogram shows almost perfect match of the field estimated with two methods across subcortical structures.

Exploration of the estimated B_1_^+^ field in sTx-MPM and pTx-MPM protocols demonstrated (Figure 7 A) an average 14% higher B_1_^+^ field in subcortical nuclei and a more consistent field settings across subjects in the pTx-MPM protocol (across subject CoV for mean field values in sTx-MPM is 0.062 versus 0.049 in pTx). In addition (Figure 7 B), pTx-MPM protocol showed more homogeneous fields across subcortical structures, demonstrating lower CoV across subcortical structures of 0.093 versus 0.118 in sTx-MPM (in group average terms).

## Discussion

We assessed how reproducible are the UHF qMRI estimates of tissue microstructural parameters of small brain structures across sites and acquisition protocols. We used age-related changes in qMRI parameters as a proxy for identifying protocol dependence of two qMRI metrics (R1 and R2*), and volume of 14 subcortical structures segmented based on multiparametric qMRI data with the same automatic approach. While the subcortical structures benefit from the increased image quality in UHF MRI their imaging might be affected by the lower reception sensitivity compared to the cortical parts of the brain. We found that for each of the quantitative parameters the age-related inter-individual variability can be correctly estimated across sites and protocols using qMRI approaches. We further observe larger relative difference across protocols for R1 and volume, while R2* remains more consistent for most regions, both in terms of difference against the reference AHEAD data (Figure 6), and in terms of displaying less interaction terms and lower amount of protocol-significant offsets.

Overall, the analysis of age-related changes demonstrates all the qMRI metrics to exhibit the inverted U-shape dependence, with the linear term in the myelin-related qMRI metrics (i.e., R1) and the iron-related metrics (i.e., R2*) showing increase with age. This is in line with the reference dataset analysis (Miletić et al., 2022) and is related to changes in brain tissue structure (e.g., myelin loss, and iron accumulation)(Bartzokis, 2011). The low number of identified interaction terms (allowing for detection power effects) between age-related variables and protocol category suggests that the age dependence matches across protocols. This is particularly true for R2* and volume measurements, having the least number of significant interaction-containing models, and less so for the R1 measurements comprising the majority of detected interactions. Nevertheless, the cohorts’ age distributions used to estimate age dependence both in MPM and AHEAD samples is uneven with fewer participants in the middle age range (30 to 50 years) suggesting the age effects might have been modelled suboptimally.

Examining the variation of qMRI metrics between MP2RAGE and variable flip angle approaches have indicated (Figure 6 A) higher R1 in the reference AHEAD data compared to the sTx-MPM and pTx-MPM data. The difference is striking when compared to the expected adult lifespan variation of the R1, showing that the estimated R1 is more affected by the protocol choice rather than participant differences. It could be speculated that due to R1 measurements in MPM protocol being strongly dependent on the knowledge of the flip angles, which is usually addressed by incorporating the field mapping (Tabelow et al., 2019), it is the field mappings (Lutti et al., 2012) that could have caused the systematic difference between the sTx-MPM and pTx-MPM results. Nevertheless, the field measurements (Figure 7 C) show that the different B1^+^ mapping methods used in sTx-MPM and pTx-MPM produce consistent estimates of the B1^+^ field. The estimated B1^+^ field (Figure 7 A) is nevertheless different in sTx-MPM and pTx-MPM configuration, suggesting it could serve as the source of the error in the R1 estimation not via the biases in the field mapping procedure, but rather due to incomplete correction of the B_1_^+^ effect during R1 calculation. This suggests further investigations into the assumptions and approximations used in the qMRI parameter extraction tools are required.

Alternatively, the discrepancy between AHEAD (MPRAGE-like acquisition) and MPMs may be induced by inherent simplification used in R1 calculation, assuming one universal R1 parameter to be able to characterize complex brain tissue, comprising multiple cellular and subcellular compartments, all exhibiting potentially different microstructural properties and macromolecular composition (Weiskopf et al., 2021). More realistic four pool models may address these simplifications, but cannot be reliably estimated from typical in vivo MRI datasets.

The R2* data exhibits (Figure 6 B) systematic differences between protocols, that are smaller than the estimated changes over the adult lifespan. This is potentially due to the less pronounced dependence of R2* on the B1^+^ field, particularly in areas with low myelin content, that results in the least amount of biases and interactions in the R2* part of the pooled dataset. The differences in estimated R2* between protocols may partly result from the model being simplified by assuming single R2* values for complex biological tissue with different compartments (Milotta et al., 2023). This results in estimated R2* depending on sequence parameters (e.g. flip angle in MPM acquisition). While these effects are expected to be most prominent in white matter, they may also affect subcortical nuclei, which are partially myelinated.

Finally, volume estimation is systematically higher in the AHEAD data (Figure 5 C), which may be related to the absence of χ maps in the MPM data and the absence of PD maps in the AHEAD data so that MASSP segmentation procedure could not be matched across all the cohorts. Notably, the difference in estimated volume between sTx-MPM and pTx-MPM protocols (Figure 6 C) is hardly significant, supporting the hypothesis of the lack of QSM data input into the segmentation algorithm being the reason for differences in the measured volume, and emphasizing the need in maximum harmonization of processing pipelines. Note that the systematic differences in segmentation procedure may have also contributed to the observed differences in qMRI values, as slightly different regions were compared across protocols.

To our knowledge this work demonstrated the first assessment of acquisition protocol effects when combining ultra-high resolution UHF qMRI data for subcortical structures. It was performed over a relatively large participant cohort, comprising data from different sites collected with different quantitative protocols. This also included assessing the impact of parallel transmission technology on the estimated quantitative parameters. Moreover, this study demonstrated the diversity of age-related and scanner-related effects across different structures by concentrating on numerous and heterogeneous subcortical structures, while previous research was mostly focused on larger tissue classes, such as white and gray matter (Al-Radaideh et al., 2015; Carradus et al., 2020; Heij et al., 2024; Piredda et al., 2023).

Examining the UHF qMRI measurements across a number of protocols and measurement sites helped identify the issues arising when attempting to merge heterogeneous datasets: the lack of consistency in absolute values of qMRI metrics, as well as the discrepancies in segmentation relying on different input data. Overall, our results suggest that while inferring age-related changes (i.e., forward inference) in e.g. iron accumulation (based on R2* values) can be reliable using pooled heterogenous qMRI datasets, inferences on the brain status, e.g., for calculating the brain age from the qMRI values (Bacas et al., 2023), or for calculating the pathology cutoff values (i.e., reverse inference (Bzdok & Ioannidis, 2019)), may be obscured by the protocol-related biases and caution should be applied. Thus, pooling across sites and protocol requires tailored harmonization procedures (e.g., ComBat) (Johnson et al., 2007; Pomponio et al., 2020), using traveling heads studies or dedicated phantoms, potentially in the framework of MRI metrology (Hall et al., 2025). R2* and ROI volume are seemingly more suited for such reverse inference, where the effect is present, or prediction in large and heterogenous multicentric studies due to their inter-protocol stability. This encourages the application of UHF qMRI to the disorders correlated with the change of the R2* in the brain tissues, such as Alzheimer’s disease, Parkinson’s disease as well as multiple sclerosis (Zecca et al., 2004).

## Acknowledgments

This work was conducted at the GIGA-In Vivo Imaging platform of ULiège, Belgium. We thank Nikita Beliy, Sophie Laloux, Annick Claes, Christian Degueldre, Brigitte Herbillon, Gregory Hammad, Benjamin Lauricella, Alexia Lesoinne, Pierre Maquet, John Read and Aubry Robert for their help in the different stages of the study. This work was supported by Fonds National de la Recherche Scientifique (FRS-FNRS, T.0242.19, T.0238.23 and J. 0222.20), the EU Joint Programme Neurodegenerative Disease Research (JPND) IRONSLEEP/HISTOPARK and SCAIFIELD projects, respectively – (FNRS references: PINT-MULTI R.8011.21/R.8002.25 & 8006.20); Action de Recherche Concertée – Fédération Wallonie-Bruxelles (ARC SLEEPDEM 17/27-09), Fondation Recherche Alzheimer (SAO-FRA 2019/0025 & 2022/0014), Fondation Léon Frédéricq, ULiège, European Regional Development Fund (Radiomed, Biomed-Hub, WALBIOIMAGING).

EB was supported by the Maastricht University - ULiège Imaging Valley. NM, CP, MZ, EL, SD, CG, FB and GV are/were supported by the FRS-FNRS. PT and LL are/were supported by the EU Joint Programme Neurodegenerative Disease Research (JPND) IRONSLEEP/HISTOPARK and SCAIFIELD projects, respectively – FNRS references: PINT-MULTI R.8011.21/R.8002.25 & 8006.20. LL is supported by the European Regional Development Fund (WALBIOIMAGING).

## Data and Code Availability

Code will be made available on request. Data is available on request, and will be made available through data sharing services.

## Author Contributions

M. Z.: Conceptualization, Methodology, Software, Formal analysis, Investigation, Data Curation, Writing - Original Draft, Writing - Review & Editing, Visualization.

K. P.: Methodology, Software, Writing - Review & Editing

P.-L. B.: Methodology, Software, Validation, Writing - Review & Editing

P. T.: Formal analysis, Investigation, Writing - Review & Editing

N. M.: Investigation

S. D.: Validation, Investigation

C. G.: Investigation

E. B.: Investigation

L. L.: Resources, Writing - Review & Editing

C. P.: Resources, Formal analysis, Writing - Review & Editing

F. C.: Resources, Writing - Review & Editing, Funding acquisition

P. M.: Validation, Investigation

E. L.: Investigation

A. A.: Methodology, Software, Writing - Review & Editing, Project administration, Funding acquisition

N. W.: Methodology, Writing - Review & Editing, Supervision, Project administration, Funding acquisition

G. V.: Conceptualization, Methodology, Validation, Writing - Review & Editing, Supervision, Project administration, Funding acquisition

E. K.: Conceptualization, Methodology, Software, Validation, Writing - Review & Editing, Supervision

