## Supplementary figures for "Pooling quantitative MRI data: A multi-protocol study of healthy subcortical ageing"

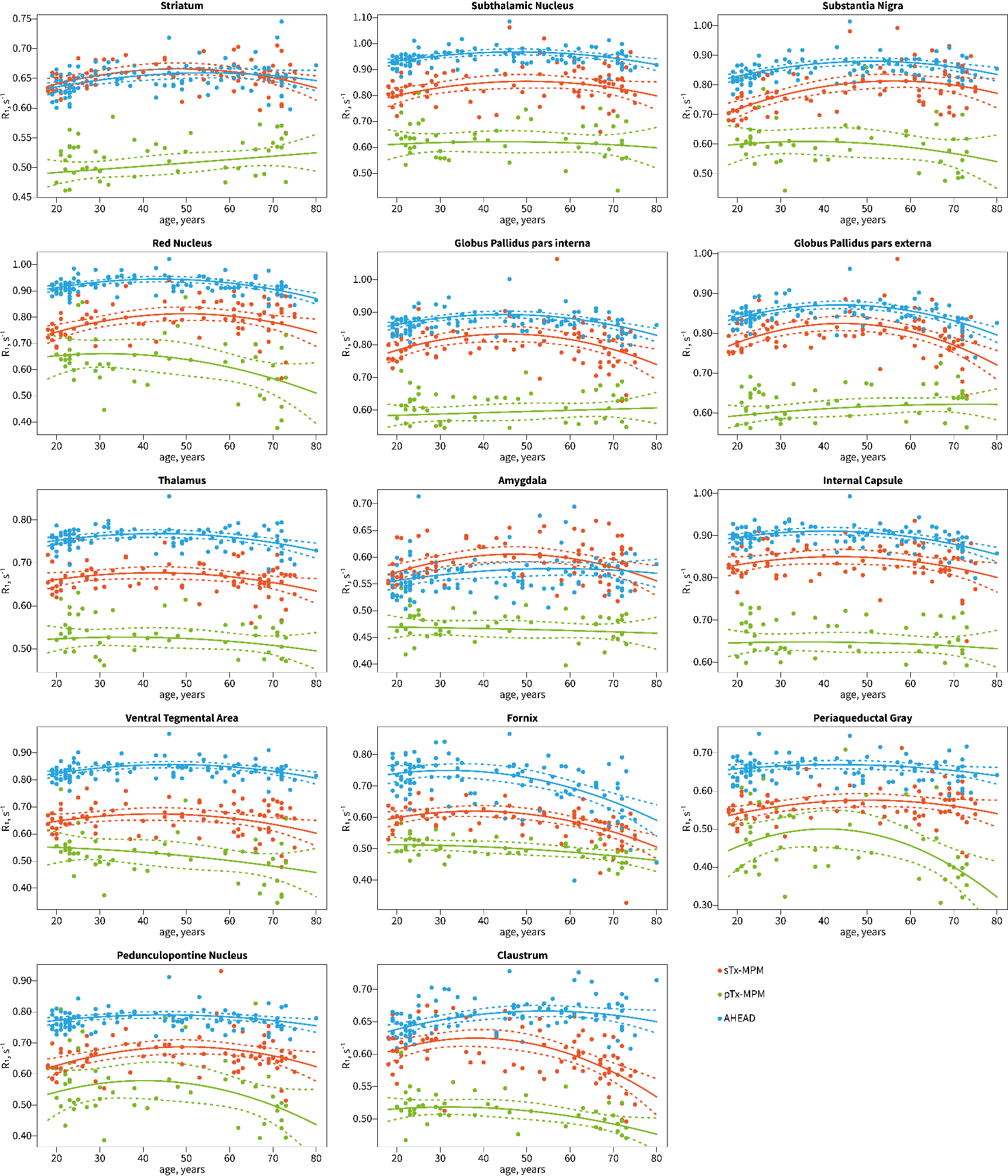

Figure S1 Median R1 values measured in the 14 subcortical ROI. Each graph shows age dependence models that were fit for each of the protocol types.

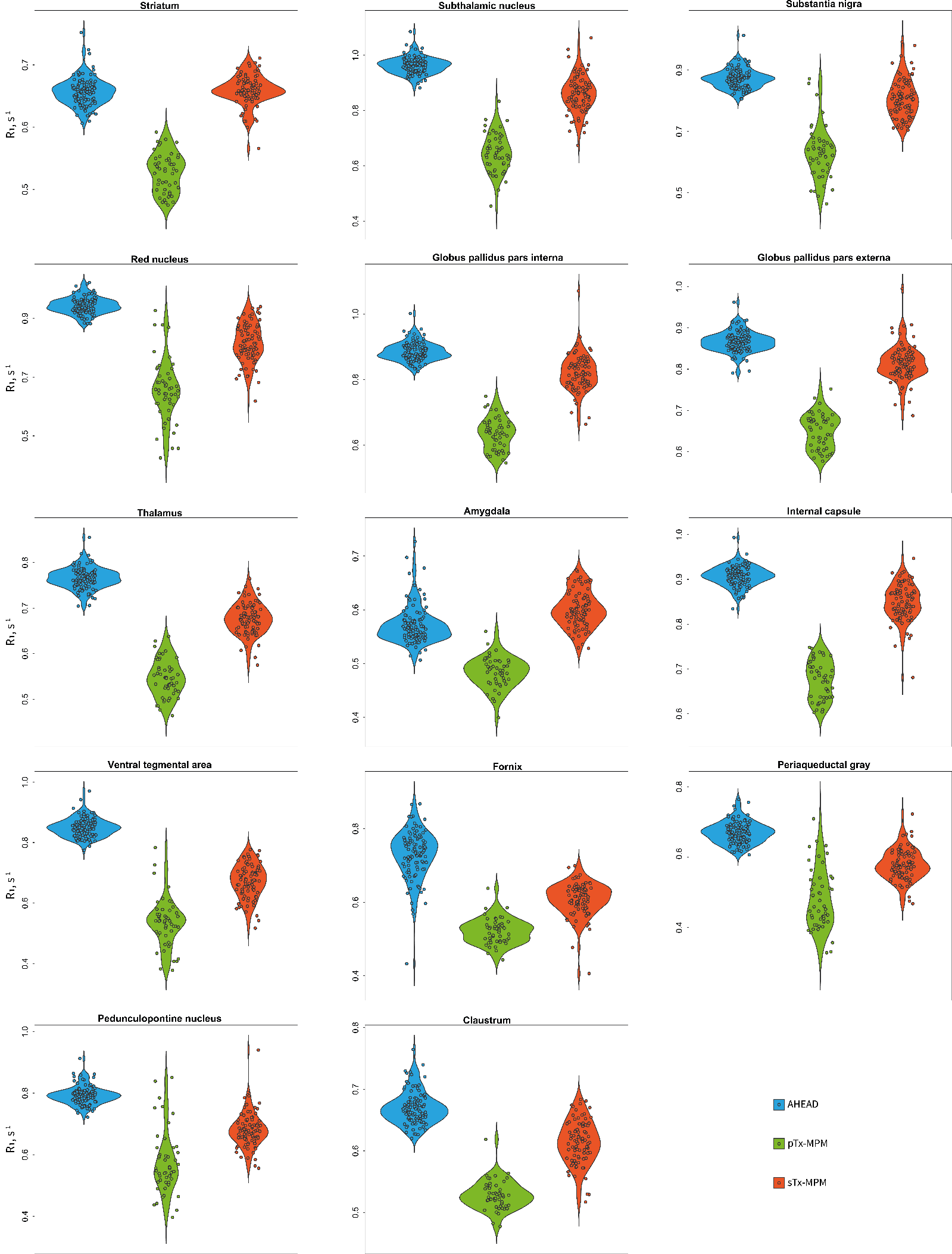

Figure S2 Protocol-stratified median R1 values measured in the 14 subcortical ROI. Graphs show data age-adjusted to the pooled dataset median age (45 years).

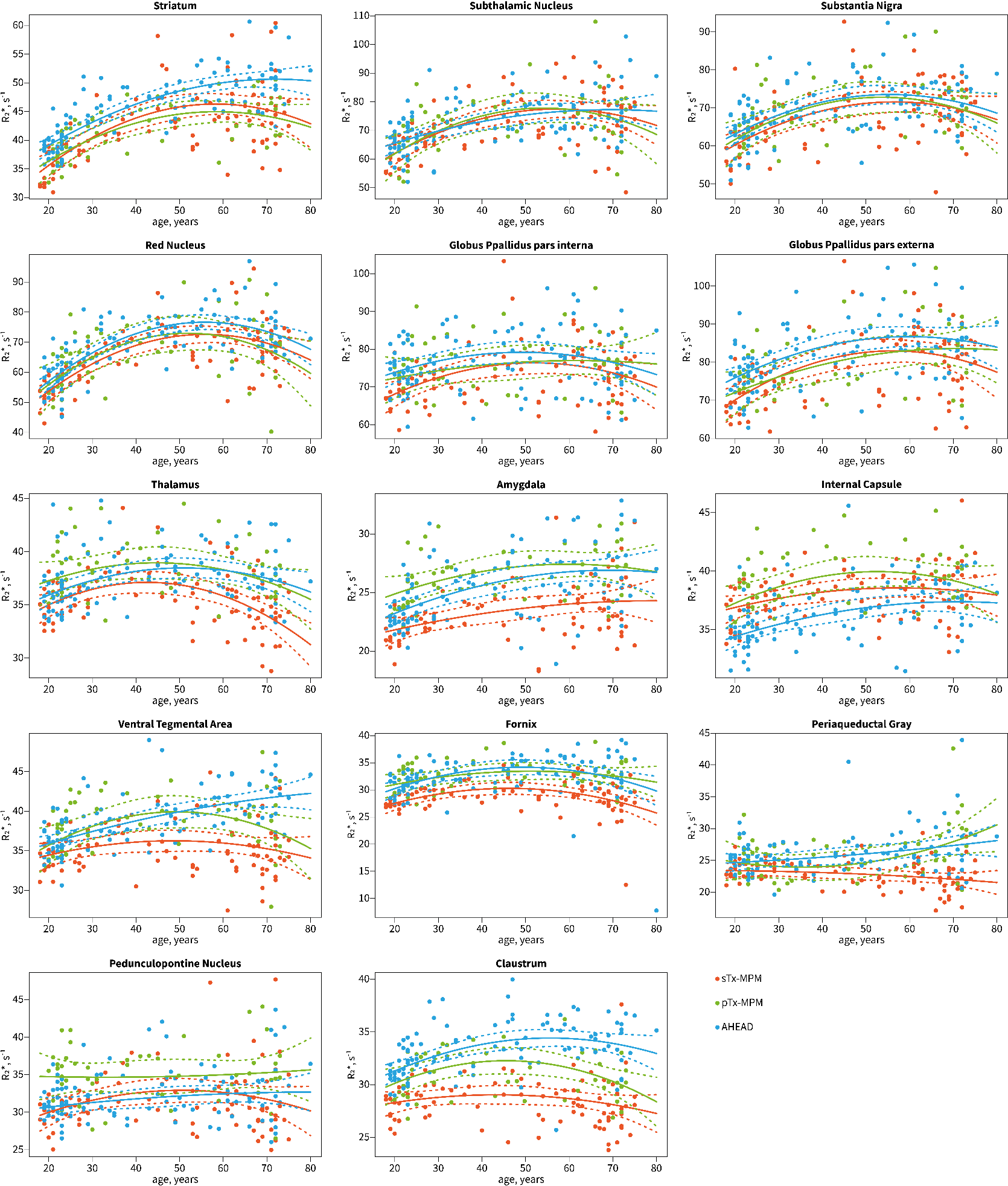

Figure S3 Median R2* values measured in the 14 subcortical ROI. Each graph shows age dependence models that were fit for each of the protocol types.

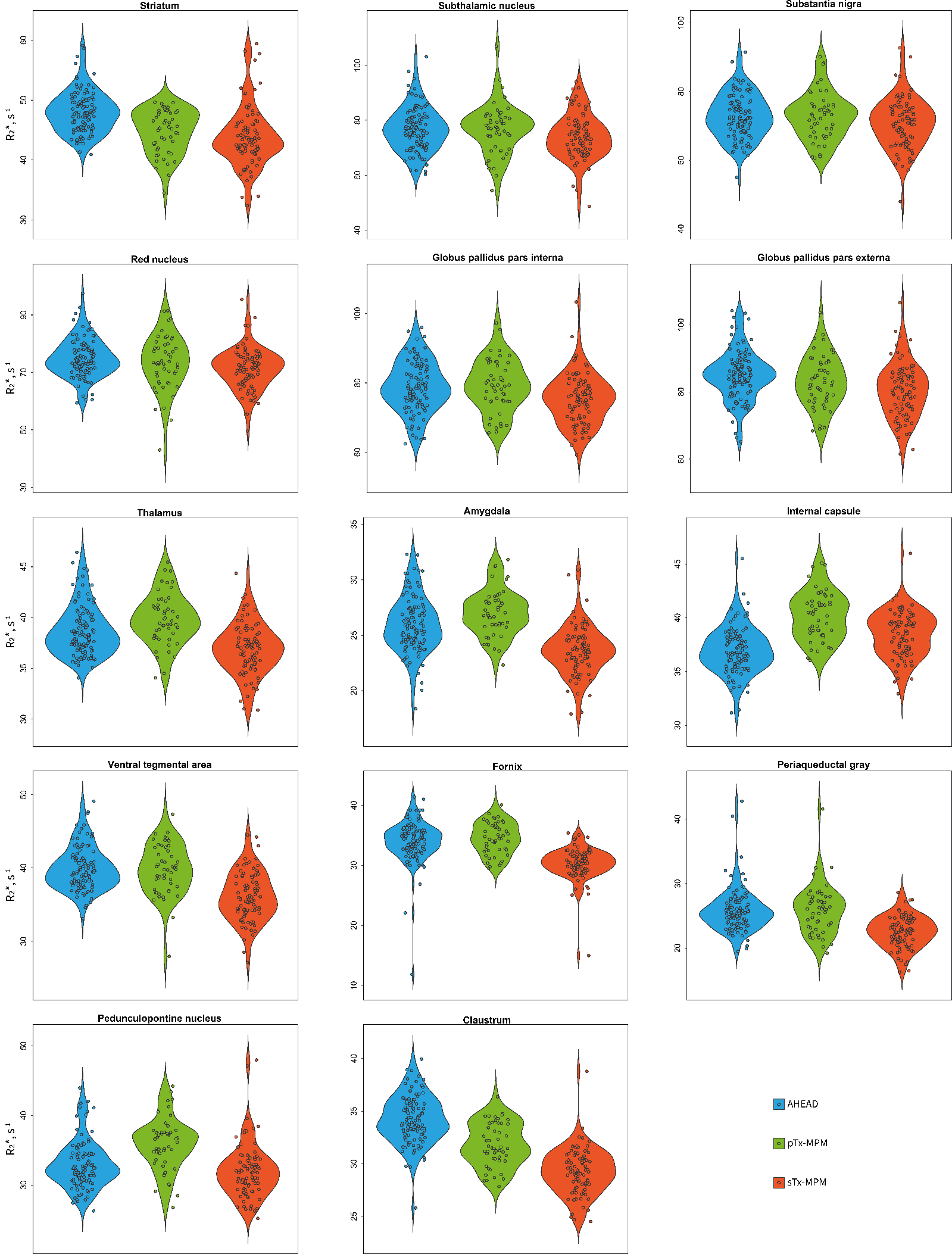

Figure S4 Protocol-stratified median R2* values measured in the 14 subcortical ROI. Graphs show data age-adjusted to the pooled dataset median age (45 years).

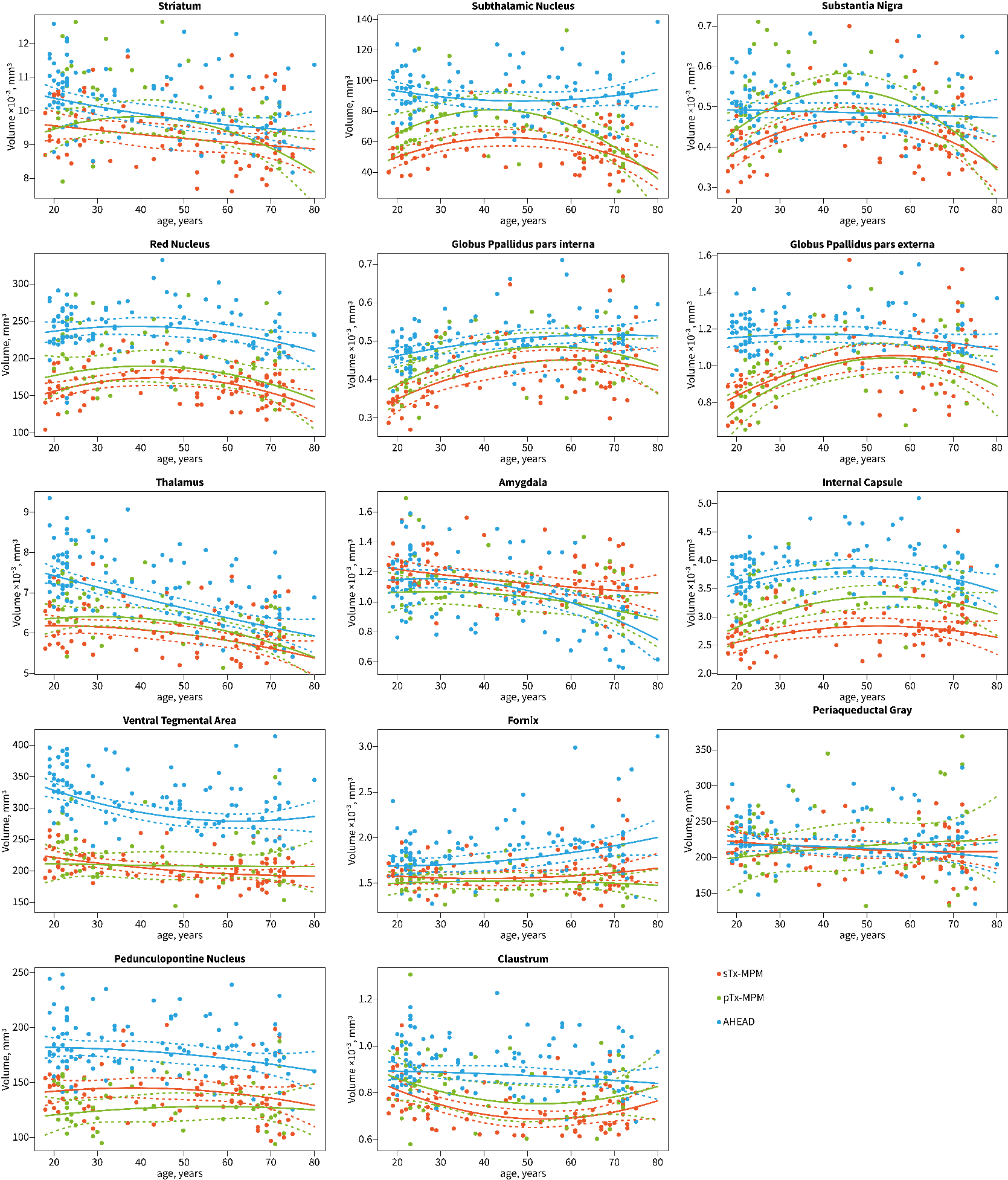

Figure S5 Volume measurements of the 14 subcortical ROI. Each graph shows age dependence models that were fit for each of the protocol types.

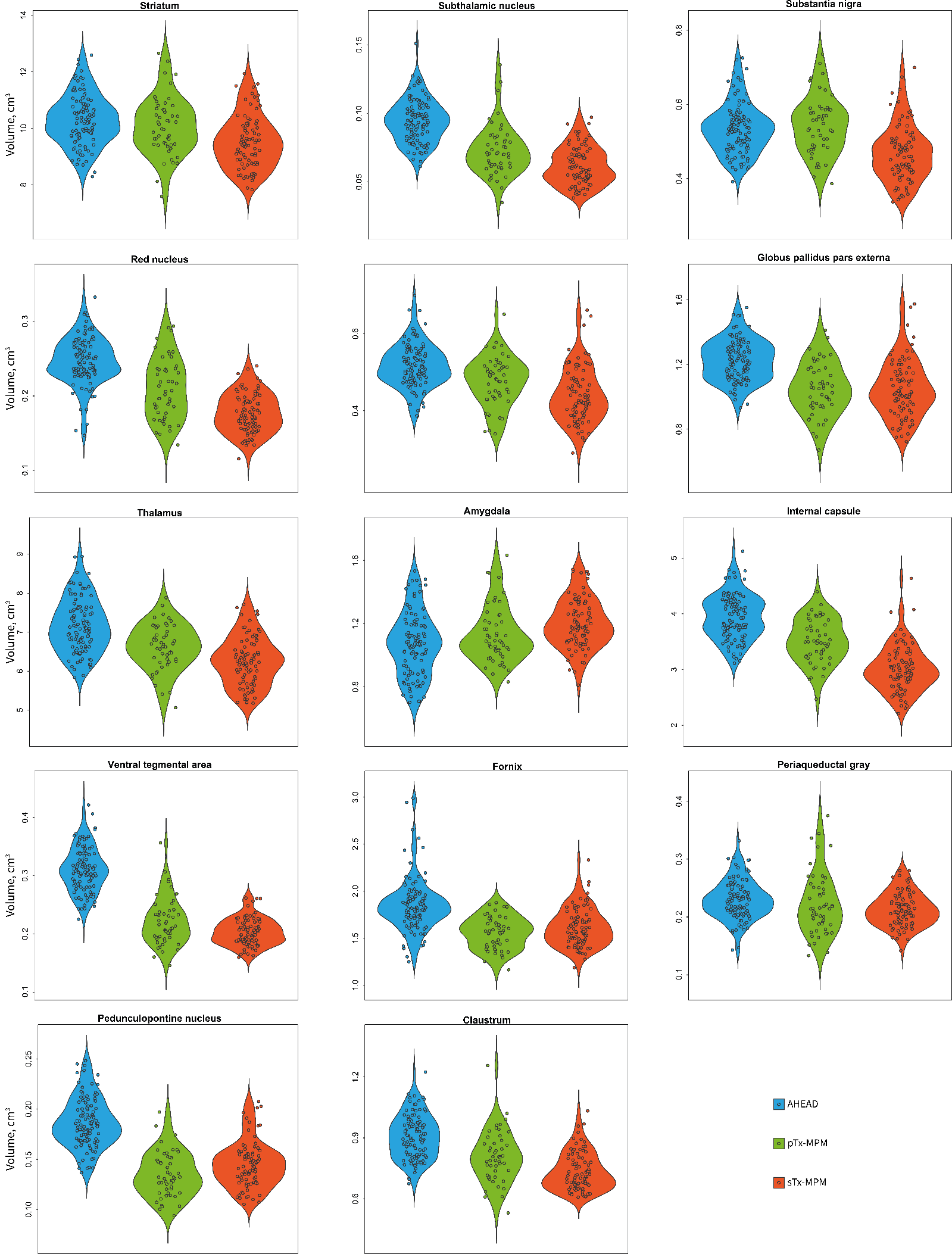

Figure S6 Protocol-stratified volume measurements of the 14 subcortical ROI. Graphs show data age-adjusted to the pooled dataset median age (45 years).

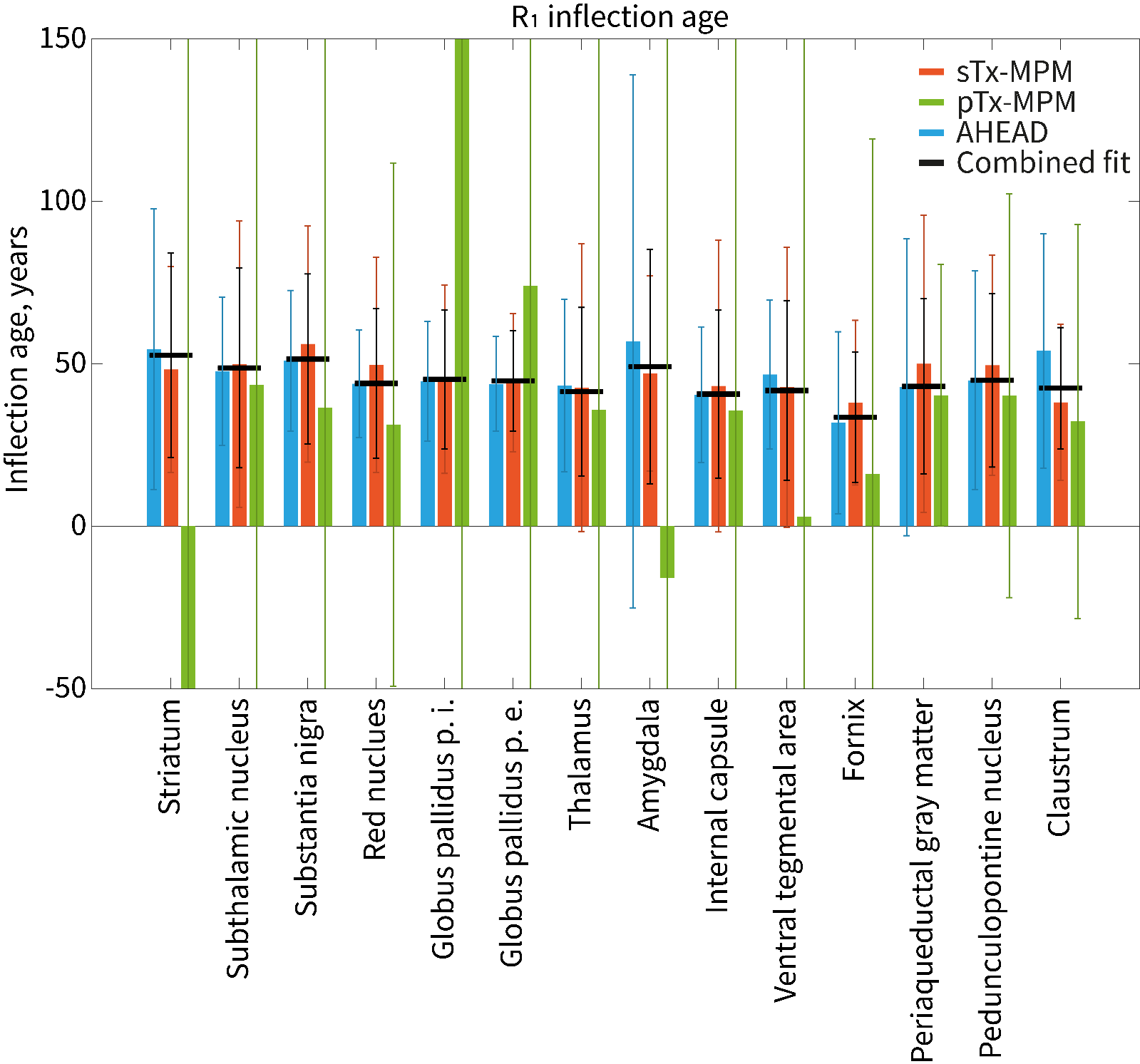

Figure S7 Ages of maximal R1 calculated from the age trajectories. The error in determining the inflection age is extremely high for each individual dataset (particularly for pTx-MPM data, exceeding often any reasonable lifespan). Note, that combining datasets does not always lead to error reduction.
